# A compact head-mounted endoscope for in vivo calcium imaging in freely-behaving mice

**DOI:** 10.1101/252205

**Authors:** Alexander D. Jacob, Adam I. Ramsaran, Andrew J. Mocle, Lina M. Tran, Chen Yan, Paul W. Frankland, Sheena A Josselyn

## Abstract

Miniaturized fluorescence microscopes for imaging calcium transients are a promising tool for investigating the relationship between behaviour and population-level neuronal activity in rodents. However, commercially available miniature microscopes may be costly, and, because they are closed-source, may not be easily modified based on particular experimental requirements. Here, we describe how to build and use a low-cost <u>c</u>ompact <u>h</u>ead-mounted <u>e</u>ndoscope (CHEndoscope) system for *in vivo* calcium imaging. The CHEndoscope uses an implanted gradient index (GRIN) lens along with the genetically encoded calcium indicator GCaMP6 to image calcium transients from hundreds of neurons simultaneously in awake behaving mice. This system is affordable, open-source, and flexible, permitting modification depending on the particular experiment. This Unit describes in detail the assembly, surgical implantation, data collection, and processing of calcium signals using the CHEndoscope system. The aim of this open framework model is to provide an accessible set of miniaturized calcium imaging tools for the neuroscience research community.

**Significance Statement:** The ability to image calcium transients in awake, behaving rodents using miniature microscopes opens exciting and novel avenues for gaining insights into how information is encoded in neural circuits. The development of this tool has already had a significant impact on neuroscience research. The cost of commercial systems, however, may be prohibitive for many laboratories. Here, we describe an affordable, open-source <u>c</u>ompact <u>h</u>ead-mounted <u>e</u>ndoscope (CHEndoscope) system for performing *in vivo* calcium imaging in freely-behaving mice. CHEndoscopes may be manufactured by individual laboratories at relatively minor cost. Our hope is that greater availability of affordable, open-source tools (such as the one presented here) will accelerate the pace of discoveries in systems neuroscience.

## Introduction

Over the past decade, the advent of novel optical technologies has revolutionized our understanding of the link between neural activity and behaviour. As the ability to target and manipulate genetically-defined populations of neurons using engineered receptors and opsins has improved, so too has the need to visualize in real-time the activity of these populations. Capitalizing on advances in protein engineering of fluorescent indicators, *in vivo* calcium imaging is a recent technique that allows neuroscientists to image the calcium activity of large populations of neurons during awake behaviour across days, weeks, or even months. This technique offers new avenues for understanding how behavioural experiences may be encoded within neural circuits.

*In vivo* calcium imaging in rodents typically relies on the expression of a genetically encoded calcium indicator (GECI) to observe neuronal activity. Of the many GECIs, GCaMP6 variants have become the standard in contemporary neuroscience studies and have been successfully employed to image neural activity in a variety of superficial and deep brain structures (Hamel, Grewe, Parker, & Schnitzer, 2015; Resendez et al., 2016). Briefly, the GCaMP protein is composed of a calmodulin domain fused to a circularly permuted enhanced green fluorescent protein moiety (Akerboom et al., 2009). Upon binding of calcium to the calmodulin domain, the protein undergoes a conformational change, shifting from a weakly-fluorescent to strongly-fluorescent state. When expressed in neurons, GCaMP exhibits increased fluorescence in response to the influx of calcium ions that accompany action potentials, and this change in fluorescence can be recorded and used as a proxy for neuronal activity.

Imaging GCaMP fluorescence in behaving animals requires optical access to the brain. Several such systems exist, including two-photon (2P) microscopy and fiber photometry. Until very recently, 2P imaging in awake mice was typically limited to surface (cortical and hippocampal) structures through a cranial window or thinned-skull preparation (Carrillo-Reid, Yang, Bando, Peterka, & Yuste, 2016; Danielson et al., 2016; Peters, Chen, & Komiyama, 2014), but recent technological advances now allow 2P imaging of deep brain structures (Otis et al., 2017). However, 2P imaging of large populations of neurons during behaviour requires head-fixation of experimental subjects, which constrains the behavioural repertoire that can be studied. In contrast, fiber photometry permits optical imaging of neural activity in unrestrained rodents but does not provide cellular resolution (Gunaydin et al., 2014).

Implantable miniature microscopes offer a means to overcome these limitations and image neuronal activity at single-cell resolution in freely behaving mice (Ghosh et al., 2011; Hamel et al., 2015; Cai et al., 2016). The microendoscope system is comprised of a gradient index (GRIN) lens which is implanted above the brain region to be imaged, along with a miniaturized epifluorescence microscope which sits atop the skull and collects images from the tissue through the implanted GRIN lens. This microscope body contains an excitation light source, optical elements which filter and focus excitation and emission light, and a detector which captures emitted fluorescence.

This Unit describes the construction and use of our recently developed <u>c</u>ompact <u>h</u>ead-mounted <u>e</u>ndoscope (CHEndoscope). The CHEndoscope is compatible with a number of established behavioural neuroscience paradigms, including, but not limited to, fear conditioning and open field exploration. The low cost of the system and flexible design allow for customization to meet specific experimental needs. In addition to construction and use, this Unit also describes a pipeline for processing calcium transient data collected using the system, including motion correction, extraction of individual calcium traces from video data, and their registration between recording sessions. Additional resources, including the design files, acquisition software, and analysis code for the CHEndoscope, are available at https://www.github.com/jf-lab/chendoscope.

## Basic Protocol 1: Assembling the CHEndoscope

The CHEndoscope system consists of three main components: the baseplate, filter box and camera body. The baseplate is implanted into the brain, and serves as a mount for the GRIN lens as well as a point of attachment for the rest of the CHEndoscope system. The filter box attaches to the baseplate, and contains an illumination LED, filters, mirrors, and lenses necessary to collect and magnify the neuronal fluorescence signal. The camera body attaches to the filter box, and houses the CMOS camera that captures imaging information relayed from the filter box. These components must be assembled prior to beginning an *in vivo* imaging experiment.

### Materials

#### Filter box

- 2.5-mm mono audio jack (Digikey, CP1-2502-ND)
- File or sanding disc
- Filter box housing (custom 3D printed)
- Two-part epoxy
- Soldering iron
- Female two-pin connector (Digikey, WM1742-ND)
- Luxeon Rebel blue LED (Digikey, 1416-1028-1-ND)
- Non-marring tweezers
- 2.4-mm Diameter × 3.0-mm Length, N-BK7 Drum Lens (Edmund Optics, 45549)
- Excitation filter (Chroma, ET470/40x)
- Dichroic mirror (Chroma, T495lpxr)
- Emission filter (Chroma, ET525/50m)
- 5-mm Dia. × 15-mm FL, VIS 0° Coated, Achromatic Lens (Edmund Optics, 49277)
- Norland optical adhesive #61 (Norland Products, NOA-61)
- 365 nm high-power UV curing light
- Superglue
- 3D printed filter box cover

#### LED power cable

- Pre-crimped cable lead (Digikey, WM2320-ND)
- Wire strippers
- Male two-pin connector (Digikey, WM1720-ND)
- Shrink wrap tubing
- 2× 1 m length hook-up wire (Digikey, 422010 BK005-ND or equivalent)
- BNC female connector with screw termination (Mouser Electronics, 992-BNC-F-TERM or equivalent)

#### Baseplate

- Baseplate housing (custom 3D printed)
- M4 × 0.5 nut (Mouser Electronics, 490-2.5MM-NUT-E)
- Two-part epoxy
- Superglue
- Activated charcoal powder
- 2 mm GRIN lens (GoFoton, ILW-200-P0250-055-NC)

#### Camera interface

- 20-pin to 5-pin connector PCB (custom ordered)
- Hot plate
- Male 20-pin connector (Digikey H121934CT-ND)
- Solder paste
- 1 m length of lightweight, four-core cable
- USB plug connector (Digikey, A107359-ND)

#### Camera body

- M8 × 1.0 tap
- M2 × 0.4 tap
- Camera body housing (custom 3D printed)
- CMOS micro-camera (Ximea, MU9PC-MBRD)
- 1.0 mm stainless steel hex nut (Scale Hardware, HXN-10-S)
- 1.0 mm stainless steel hex bolt (Scale Hardware, HXB-10-3-S)
- Two M2.5 × 6-mm nylon screws (Mouser Electronics, 534-29331)
- M2 × 4-mm nylon screw (Duratool, DTRNSE 1207 M2 4)

### Assemble the filter box

1. Remove the metal M4 × 0.5 thread bolt from the 2.5-mm audio jack. Using a file or sanding disc, sand the length of the bolt from 3.5mm down to 2mm.
2. Attach the filed M4 × 0.5 thread bolt to the base of the printed filter box housing using two-part epoxy (see Figure 1A, part no. 7). Let epoxy set for 24 h.
3. Solder the connection points of the female two-pin connector to the positive and negative solder pads of the Luxeon Rebel LED.
4. Using non-marring tweezers, insert the 2.4-mm diameter drum lens into the filter box housing as illustrated in Figure 1A.
5. Position the LED with the half-ball lens facing toward the filter box housing. Fix in place with two-part epoxy and let set for 24 h.
6. Insert the excitation filter, emission filter and dichroic mirror into their respective positions using non-marring tweezers (Figure 1A, part no. 8-10).
7. Place two small drops of Norland optical adhesive into top threaded chamber of the filter box housing, then gently insert the 5-mm diameter achromatic doublet lens using non-marring tweezers. Cure the optical adhesive with 365 nm UV light to secure the achromatic lens in place. *Ensure that the achromatic doublet lens is in the correct orientation (more convex face pointing downward) before securing in place.*
8. Place a small drop of superglue on the inset plastic region to the left of the emission filter. Press the filter box cover so it fits snugly over the chamber to seal the filter box.

**Figure 1.**
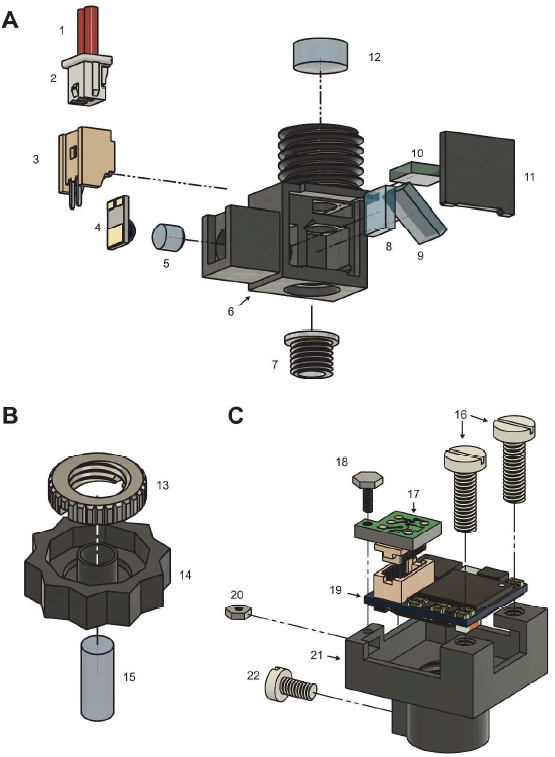
Assembly of the CHEndoscope. (A) Exploded view of the filter box with numbered components. (B) Exploded view of the baseplate. (C) Exploded view of the camera body. Numbered parts: (1) Crimped cable lead (2) Male two-pin connector (3) Female two-pin connector (4) LED (5) 2.4-mm diameter drum lens (6) filter box housing (7) M4 × 0.5 thread bolt (8) excitation filter (9) dichroic mirror (10) emission filter (11) filter box cover (12) achromatic doublet lens (13) M4 × 0.5 nut (14) baseplate housing (15) 2-mm diameter GRIN lens (16) M2.5 × 6mm nylon screw (17) camera interface PCB (18) 1-mm length hex bolt (19) CMOS camera board (20) 1-mm length hex nut (21) camera body housing (22) M2 × 4-mm nylon screw 274x289mm (300 × 300 DPI)

### Construct the LED power cable

9. Cut a pre-crimped cable lead in half and strip a small length of the cut ends, rendering two equally sized cable segments. *Steps 9-12 detail how to assemble the LED power cable without a crimper. However, if a crimper (E.g. Digikey, WM15815-ND) is available, these steps can be replaced by crimping a connection terminal (Digikey, WM1142CT-ND) directly onto a 1-m length of hook-up wire, and inserting the crimped ends into the male two-pin connector)*.
10. Insert the crimped ends of each cable segment into the male two-pin connector Ensure the crimped leads are inserted in the correct orientation, and that the connection is secure before proceeding.
11. Cut two 1-m lengths of hook-up wire and strip a small portion of each end.
12. Solder one end of each length of hook-up wire to the stripped end of each cable segment. Secure the soldered joints by enclosing them in heat-shrink tubing.
13. Insert the remaining ends of the hook-up wire into the positive and negative screw terminals of the BNC connector. *The BNC connector used in this step can be substituted for any number of other connector terminations compatible with a DC power source of your choice.*
14. Insert the male two-pin connector into the female two-pin connector on the CHEndoscope filter box to complete the LED power supply. *The connection between the LED and power cable is detachable, allowing for unhindered rotation of the camera body during large focus changes.*

### Assemble the baseplate

15. Line the interior flat surface of a 3D printed baseplate housing with two-part epoxy. Insert an M4 × 0.5 nut into this pocket and secure in place with additional epoxy. Let epoxy set for 24 h.
16. Invert the baseplate housing so that the nut is facing downward. Place on a leveled, flat surface
17. Prepare an opaque superglue solution by mixing a small drop of superglue with activated charcoal.
18. Using non-marring tweezers, insert a 2-mm GRIN lens into the central bore of the baseplate housing. Ensure the lens is positioned perpendicular to the flat surface of the baseplate housing, then secure the lens in place with a small quantity of opaque superglue. Let superglue set for at least 4 h. *It is important that the GRIN lens sits perpendicular to the bottom of the baseplate. It may be necessary to adjust the position of the lens by gently pressing on the side of the lens with a pair of non-marring tweezers once the superglue has been applied.*

### Construct the camera interface PCB

19. Place a custom 20-pin to 5-pin PCB on a hot plate (powered off) with CMOS interface side (Figure 2A) facing up. *Note: Surface mounted components are ideally soldered using a reflow oven, however this equipment is not commonly available in neuroscience laboratories. Here we describe how the interface PCB is soldered to the male 20-pin connector using a hot plate, a tool which is more widely available.*
20. Using forceps, place a male 20-pin connector onto the custom PCB such that the metal terminals of the connector are aligned with the pads of the PCB (Figure 2B).
21. Apply solder paste separately to each of the 20 metal terminals. *It is important that no bridges form between adjacent terminals. To ensure this, apply solder paste carefully using a fine-gauge needle tip. A small amount of solder paste is sufficient for each terminal*.
22. Turn on the hot plate to 300°C, and observe the solder paste as the temperature gradually rises. When the solder has melted and pooled between the terminals and pads of the PCB, carefully remove the PCB assembly from the heat. *A dissecting microscope positioned above the hot plate can be useful for monitoring the status of the solder paste. If the PCB is heated for too long, it may become damaged*.
23. Turn the PCB assembly over and identify the four solder pads on the USB interface side labelled D+, D-, 5V, and GND (Figure 2C). Place a drop of solder on each pad. *The SYNC connection listed in Figure 2C is not used in this protocol. Do not solder this pad*.
24. Strip both ends of a 1-m length of four-core cable. At one end of the cable, solder each of the four wires to the corresponding pads on the USB interface side of the PCB assembly. *The cabling used in this step must be lightweight and flexible to allow for unrestrained movement of the mouse. Our lab uses a cable composed of four strands of 19 × 0.04mm^2^ twisted copper wire wrapped in an unshielded 2 mm silicon rubber sleeve. We observe stable USB signals in this cable at lengths exceeding 1 m*.
25. Secure the soldered wires in place using optical adhesive. Apply a large drop to surround the four soldered connections and cable sleeve, then cure the adhesive with UV light. Several applications of adhesive may be necessary to stabilize the soldered connection.
26. At the other end of the four-strand cable, solder each of the wires to a corresponding position on the USB plug connector (Figure 2D). *Ensure that each of the coloured wires correctly connects a labelled pad on the PCB assembly to the corresponding position on the USB interface*.
27. Enclose the USB interface jack in its included metal jacket to finish the camera-to-USB cable. *Optional: the completed LED power cable (steps 12-16) can be wrapped around the USB cable*.

**Figure 2.**
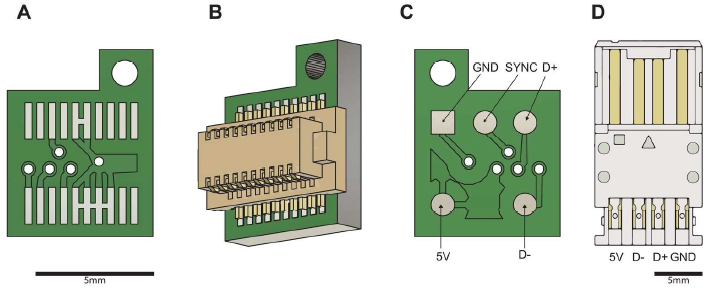
Schematic of camera-to-computer interface components. (A) CMOS-facing side of the interface PCB (B) Rendering of the interface PCB with attached male 20-pin connector (C) USB-facing side of the interface PCB. The USB cable outputs for 5 V power, GND (ground), D+ (data positive) and D- (data negative) are indicated. (D) USB plug connector. USB inputs are labelled as in (C) 180x75mm (300 × 300 DPI)

### Assemble the camera body

28. Insert the M8 × 1.0 tap into the bottom of the 3D printed camera body housing, then twist the tap through the printed threads to clear out excess plastic. Repeat this process with the smaller M4 × 0.2 tap for the set screw hole in the side of the camera body housing.
29. Connect the female 20-pin receptacle of the Ximea CMOS camera to the male 20-pin connector of the camera interface PCB *Consult Figure 1C (part no. 17) to ensure the camera and PCB are connected in the correct orientation.*
30. Seat the PCB-camera assembly in the camera body housing.
31. Align the through-hole of the camera interface PCB with the corresponding hole in the camera body housing. Place a 1-mm metal hex nut through the holes, and secure in place with a 1mm hex bolt.
32. Insert two M2.5 × 6-mm screws into the top-facing holes in the camera body housing.
33. Insert an M2 × 4-mm screw into the side of the camera body housing.

## Basic Protocol 2: Virus Infusion Surgery

The CHEndoscope system requires one or two surgical procedures, depending on the method used to express GCaMP in the brain region of interest. The purpose of these procedures is to (1) infuse an adeno-associated virus (AAV) carrying the GCaMP transgene in the brain region of interest, and (2) implant the baseplate assembly above the region of interest for later visualization of calcium activity using the CHEndoscope. This protocol describes infusion of an AAV expressing GCaMP6f into the CA1 region of the dorsal hippocampus of wild-type mice. Notably, by modifying the stereotaxic coordinates, the procedure outlined here can be used for infusion of virus into other brain regions. This protocol can be skipped if using transgenic mice that already express GCaMP in the target brain region (see Critical Parameters section for considerations on selecting viral versus transgenic approaches for expressing GCaMP).

### Materials

- Adeno-associated virus (AAV) carrying GCaMP6f transgene *e.g., AAV(DJ)-CaMK2α-GCaMP6f, obtained from Stanford University Gene and Viral Vector Core (AAV-90) or made in-house*
- P10 pipette
- Filter pipette tips
- 0.5 ml PCR tubes
- Dry ice
- 70% (v/v) ethanol
- 1-ml syringes
- 26-gauge needles
- Appropriate anesthetic
- Atropine sulfate (0.08 mg/mL in sterile water)
- Sterile saline (0.9% sodium chloride, for injection)
- Metacam (0.25 mg/mL in sterile saline)
- Eye lubricant
- Stereotaxic frame for mouse
- Stereotax arm with clamp (for holding micropipette)
- Virus aliquots
- Glass micropipette (opening cut to 20 um diameter) with attached polyethylene tubing
- 10-ul needle syringe and programmable micro-infusion pump, or other infusion system
- Distilled water
- Cotton swabs (autoclaved)
- Curved forceps
- Fine surgical scissors or scalpel
- Microdrill
- 0.5 mm burr for microdrill
- Dissolvable sutures
- Antibiotic ointment
- Clean mouse cage
- Heating pad

### Prepare virus aliquots

1. Remove stock of AAV (purchased or produced in-house) from −80 °C freezer and place stock tube over wet ice until completely thawed.
2. Aliquot the entire stock of AAV construct into 0.5 ml tubes with 4-5 µL of virus in each tube. Place each tube immediately on dry ice.
3. Disinfect all work surfaces and waste materials with 70% EtOH before discarding.
4. Store AAV aliquots at −80 °C. *With minimal freeze/thaw cycles, virus can be stored at −80 °C for several years.*

### Prepare mouse for viral infusion

5. Anesthetize mouse (5-6 weeks of age) with appropriate anesthetic and treat with atropine sulfate (0.1 mg/kg, i.p.). When fully anesthetized, place mouse in stereotaxic frame.
6. Inject 0.5 ml warm saline (s.c.) and metacam (2 mg/kg, s.c.) for hydration and analgesia during the surgery.
7. Set up the infusion system by placing the glass micropipette into the stereotax arm holder and carefully withdrawing the virus into the micropipette tip.
8. Gently wipe the head of the mouse with an ethanol-soaked cotton swab 3 times to sterilize the incision site.
9. Expose the skull by using fine scissors or a scalpel to make an incision along the midline. The incision should stop slightly above and below bregma and lambda, respectively. Ensure that the skull is leveled in both the anterior-posterior and medial-lateral axes. If it is not, make the necessary adjustments to the height of the bite bar and/or ear bars.

### Perform craniotomy and infuse AAV into the brain

10. Measure bregma by manipulating the stereotax arm to place the tip of the micropipette at bregma. Record bregma coordinates (AP, ML, and DV).
11. Raise the micropipette off the skull surface and move the micropipette to the injection coordinates (for CA1 region of dorsal hippocampus, −1.9 mm AP and +1.5 mm ML from bregma).
12. Insert the burr into the microdrill and make the craniotomy immediately below the micropipette tip. Gently use cotton swabs to clear any debris around the craniotomy.
13. Slowly lower the micropipette into the brain to the depth of the CA1 pyramidal layer (-1.5 mm DV from bregma), start the infusion pump, and wait for the infusion to finish (5 min; 0.5 µL at 0.1 µL /min). Visually confirm that the infusion was successful by checking that volume of virus in the micropipette has decreased.
14. Leave the micropipette in the brain for additional 10 min from the end of the virus infusion to allow for diffusion of the AAV into the tissue. *Removing the pipette immediately following the infusion can result in reflux of the AAV and low expression levels.*
15. Slowly raise the micropipette to remove it from the brain, and remove the mouse from the stereotax.

### Close the incision for recovery

16. Clean the incision with cotton swabs, then close the incision with 2-3 sutures.
17. Apply antibiotic ointment to the closed incision using a cotton swab.
18. Inject an additional 0.5 ml warm saline subcutaneously and place the mouse in a clean cage on a heating pad until fully recovered from anaesthesia.
19. Perform daily post-operative monitoring for at least 3 days following the surgery, and administer analgesic and fluids PRN.
20. Allow mice to recover for at least 1 week before GRIN lens implantation.

## Basic Protocol 3: Lens Implantation Surgery

In order to visualize neuronal activity using the CHEndoscope, a GRIN lens must be implanted above the target brain region and a baseplate installed around the lens on the head of the mouse to interface with the microscope. Unlike previous microendoscope systems that require separate procedures for lens implantation and installation (Cai et al., 2016; Grewe et al., 2017; Resendez et al., 2016; Ziv et al., 2013), in our system, the GRIN lens and baseplate are pre-attached and implanted into the brain in a single process. The implantation surgery is identical whether using a viral or transgenic strategy to express GCaMP in the cell population of interest.

### Materials

- Pre-assembled baseplate with GRIN lens (see Basic Protocol 1)
- 20- and 26-gauge needles
- Blunted, 45° angled 20- and 26-gauge needles
- Artificial cerebrospinal fluid (aCSF; see Reagents)
- 1- and 3-ml syringes
- Appropriate anesthetic
- Atropine sulfate (0.08 mg/mL in sterile water)
- Dexamethasone (0.5 mg/mL in sterile saline)
- Stereotaxic frame for mouse
- Sterile saline (0.9% sodium chloride, for injection)
- Metacam (0.25 mg/mL in sterile saline)
- Eye lubricant
- Microdrill
- 0.5 mm burr for microdrill
- Stereotax arm with microdrill holder
- 70% ethanol
- Cotton swabs (autoclaved)
- Curved forceps
- Fine surgical scissors or scalpel
- Bulldog serrifines (x4)
- 3% hydrogen peroxide
- 1.19-mm diameter machine screws (PlasticsOne, 00-96 × 1/16)
- Slotted screwdriver (e.g. PlasticsOne, SD-96)
- 2.3 mm trephine for microdrill
- Dissecting microscope
- Size 5 forceps
- Vacuum pump for aspiration
- Stereotax arm with filter box holder (custom, 3D-printed)
- Filter box (custom, 3D-printed)
- Dental cement liquid and powder (black)
- Clean mouse cage
- Heating pad

### Prepare materials for implantation surgery

1. Assemble the appropriate number of baseplates fitted with GRIN lenses for implantation (see Basic Protocol 1).
2. Prepare aCSF (see Reagents) and store at 4 °C.

### Prepare mouse for baseplate implantation

3. Anesthetize mouse (6-7 weeks of age, 1 week following viral infusion if using viral strategy for GCaMP6f expression) and treat with atropine sulfate (0.1 mg/kg, i.p.). Inject mouse with dexamethasone (5 mg/kg, i.p.) to reduce inflammation and swelling during brain aspiration. When fully anesthetized, place mouse in stereotaxic frame.
4. Inject 0.5 ml warm saline (s.c.) and metacam (2 mg/kg, s.c.) for hydration and analgesia during the surgery, and apply eye lubricant.
5. Gently scrub the head with an ethanol-soaked cotton swab 3 times to sterilize the incision site.
6. Make a large incision along the midline of the scalp using fine scissors or scalpel, exposing lambda and 2-3 mm of skull anterior to bregma. If using mice previously infused with AAV, cut along the partially healed incision made previously. Make any adjustments required to level the skull both the anterior-posterior and medial-lateral axes.
7. Clamp the incision in 4 corners with bulldog serrifines. Allow the bulldog sereffines to hang from the mouse’s head to retract the skin and widen the surgical field (see Figure 3A).
8. Gently scrub the surface of the skull with a cotton swab dipped in 3% hydrogen peroxide to remove all connective and scar tissues visible on the skull surface. Dry the skull with a clean cotton swab following hydrogen peroxide application.

**Figure 3.**
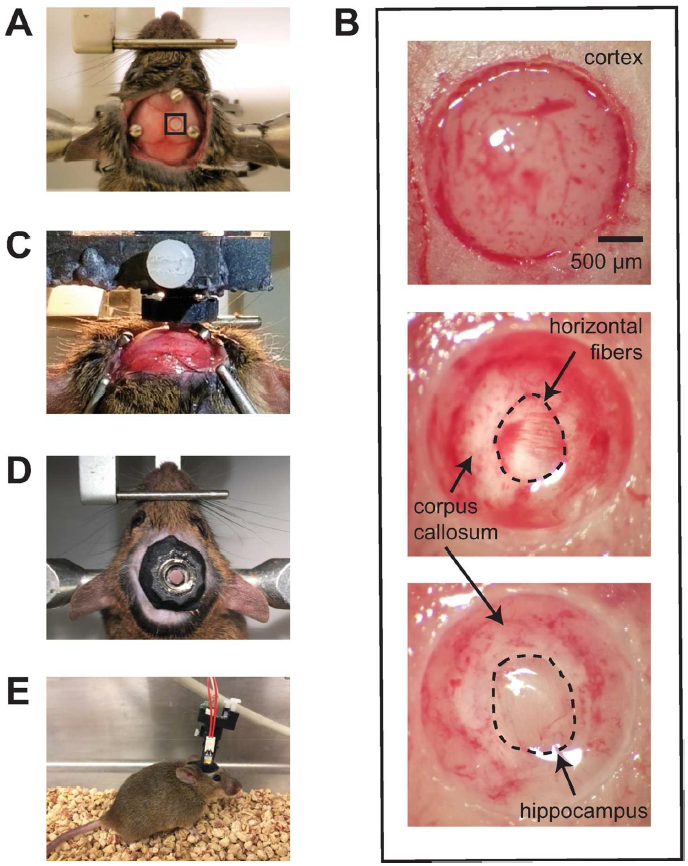
Surgical implantation of the baseplate assembly. (A) A craniotomy is made above the hippocampus using a trephine bit (black box) and screws are implanted around the site to later anchor the baseplate to the skull. (B) Magnified view of area depicted in black box Panel A. Immediately following the durotomy, the cortex is exposed (top). The cortex and corpus callosum are slowly aspirated with vacuum suction through a large, blunted needle until fibers running horizontally become visible (center). Tissue aspiration is further performed with a smaller, blunted needle to remove the horizontal fibers and reveal the hippocampal surface containing the imaging plane (bottom). (C) Following the aspiration and cessation of major bleeding, the baseplate assembly is attached to a filter box and mounted on the stereotax and the GRIN lens is lowered into the craniotomy above the hippocampus. (D) Dark dental cement is applied between the skull and baseplate using syringes to secure the baseplate in place. (E) After recovery, the CHEndoscope can be attached the baseplate and carried by the mouse during unrestrained behaviour. 112x139mm (300 × 300 DPI)

### Insert skull screws and perform craniotomy with trephine

9. Insert the microdrill into the holder on the stereotax arm and insert the burr into the drill. Attach the drill and holder to the stereotax.
10. Manipulate the stereotax arm to place the tip of the drill burr on top of bregma and record the AP and ML coordinates.
11. Raise the drill off of the skull and move the burr to the site of the craniotomy (-2.1 mm AP and +1.5 mm ML from bregma). Turn on the drill and slowly lower it towards to skull to “mark” the location of the craniotomy.
12. Detach the stereotax arm and while holding the microdrill, use the burr to drill 3 holes where screws will be placed. *Figure 3A illustrates screw placements that do not interfere with baseplate implantation above CA1 region of the dorsal hippocampus.*
13. Use curved forceps to align each screw with the drilled hole, and tighten with a screwdriver until the screw is firmly implanted in the skull. *The screws are the anchor points of the baseplate implant. It is important to ensure the stability of the screws relative to the skull. A stable screw should not display any noticeable movement with a gentle nudge.*
14. Remove the burr from the microdrill and insert the 2.3 mm trephine. Place the microdrill back on the stereotax.
15. Move the stereotax arm to center the trephine over the craniotomy site (-2.1 mm AP and +1.5 mm ML). *The previous “mark” made with the burr should be centered within the ring of the trephine.*
16. Turn on the drill and slowly lower it toward the skull. Continue to thin the skull until the center fragment separates from the remainder of the skull. Gently clear any debris from the area of the craniotomy with a cotton swab before continuing.
17. Reposition the dissecting microscope for use and magnify (4-5x magnification) to bring the craniotomy into view.

### Perform the durotomy

18. While looking through the dissecting microscope, carefully remove the detached bone fragment from over the craniotomy using size 5 forceps and gently remove any other bone fragments that may be partially attached to the skull.
19. Run the tip of the size 5 forceps around the edge of the craniotomy to break and pull back the meninges. *Breaking the meninges will fill the craniotomy with blood, which will be cleared in the following steps.*

### Perform the aspiration

20. Fill a 3-ml syringe with sterile aCSF and attach a blunted 26-gauge needle to the end.
21. Turn on the vacuum pump and attach a blunted and bent 20-gauge needle to the end of the aspiration tubing.
22. Hold the aCSF-filled syringe in one hand and the 20-gauge aspiration needle in the other. While dripping aCSF over the exposed tissue, begin to carefully suction the cortical tissue with the 20-gauge needle by slowly moving it in a circular motion around the inside edge of the craniotomy. *Continue to slowly drip aCSF over the tissue throughout the entirety of the aspiration procedure to aid in tissue removal. Ensure that the tissue is always covered in aCSF to prevent dehydration.*
23. Remove all cortical layers using the 20-gauge needle, then the corpus callosum. *The corpus callosum can be visually identified by a change in tissue colour. Once the cortical layers have been completely removed, the tissue will appear very pale which can be used as an indicator of the corpus callosum white matter tracts.*
24. Stop aspirating with the 20-gauge needle when dense, white fibers running horizontally (i.e., perpendicular to the sagittal sinus) become visible (see Figure 3B). Remove the 20-gauge needle tip and replace it with a blunted and bent 26-gauge needle.
25. While suctioning, move the 26-gauge needle around the edge of the craniotomy close to the tissue to break and remove the horizontal fibers above the hippocampus. *Do not bring the needle in direct contact with the tissue, but instead allow the suction to draw the fibers to the needle tip. This prevents damage to the hippocampal tissue which is immediately below this white matter layer. Visually identify the hippocampal tissue by its pink hue relative to the horizontal white matter fibers. A more sparse layer of vertical fibers (i.e., parallel to the sagittal sinus) immediately below the horizontal fibers and dorsal to the hippocampus can be left undisturbed.*
26. Continue to break the horizontal fibers and remove them using suction to create a large enough surface of hippocampal tissue on which to image. *The center of the craniotomy containing the imaging surface should be almost void of white matter tracts; however, it is permissible for some white matter to remain around the edges of the craniotomy.*
27. When the aspiration is complete, continue to irrigate the craniotomy with aCSF and suction it away until all major bleeding has ceased (approximately 5-10 min).

### Implant the baseplate

28. Screw a pre-assembled baseplate onto the end of the filter box and place the filter box into the stereotax arm with the custom holder. Attach the stereotax arm to the stereotaxic frame.
29. Manipulate the stereotax arm to bring the bottom of the GRIN lens 1-2 mm from the surface of the skull and adjust the filter box in the holder so that the GRIN lens surface is parallel with the skull surface.
30. Move the stereotax arm to center the GRIN lens above the craniotomy.
31. Slowly lower the GRIN lens until the bottom of the lens reaches the skull surface from the medial side. Record the DV coordinate.
32. Slowly lower the GRIN lens into the craniotomy to −1.5 mm below the skull surface.

### Create the headcap by cementing the baseplate into place

33. Use cotton swabs to dry the surrounding skull and skin, while avoiding contact with the baseplate.
34. Drop approximately 1 ml of dental cement liquid into a medicine cup and mix in a slightly smaller amount of dental cement powder.
35. Withdraw the mixture into a 3-ml syringe and cap with a blunted 20-gauge needle.
36. Place the tip of the needle between the baseplate and the skull and pipe the liquid cement mixture into the area such that it covers the entire surface of the exposed skull, surrounds the anchor screws, and makes contact with the underside of the baseplate.
37. Remove the bulldog serrifines from the edges of the incision.
38. Add more dental cement around all sides of the baseplate to strengthen the headcap. The height of the headcap should not exceed the height of the baseplate.
39. Allow headcap to completely dry (approx. 15-20 min).

### Detachment of the filter box and recovery

40. Loosen the filter box from the holder and carefully detach and remove the stereotax arm.
41. Unscrew the filter box from the now implanted baseplate.
42. Inject an additional 0.5 ml warm saline subcutaneously and place the mouse in a clean cage on a heating pad until it recovers from anaesthesia.
43. Perform daily post-operative monitoring for at least 3 days following the surgery, and administer analgesic and fluids PRN.

## Basic Protocol 4: Preparing for Imaging

*In vivo* imaging experiments require both high-quality recordings and robust behaviour. To this end, preparations should be made in advance of the experiments. First, after recovery from lens implantation surgery (3-4 weeks post-surgery) mice should be assessed using the CHEndoscope. This allows identification of imaging planes and settings for the imaging conducted during the experiment. Second, to reduce the impact of the CHEndoscope (e.g., attachment, weight, etc.) on the subjects’ behaviour during the experiment, mice should be habituated to microscope procedures. The habituation procedure is ideally performed in the few days preceding the start of the experiment.

### Materials

- 100% EtOH
- Cotton swabs or lens paper
- CHEndoscope
- Clean mouse cage
- Computer with Python (RRID:SCR_008394) and CHEndoscope acquisition software (found at https://github.com/jf-lab/jflab-minipipe)
- Adjustable DC power source

### Attach the CHEndoscope to the mouse

1. Remove one mouse from its home cage and gently restrain the mouse by holding the skin at the base of the baseplate and tail with one hand. *Restraining the mouse by scruff is also effective, but can stress the animal and result in resistance.*
2. While restraining the mouse, clean any dirt or debris from the top of the GRIN lens with 100% ethanol and a cotton swab or lens paper.
3. Hold the microscope at the filter box (avoid handling the microscope by the camera body and the LED) and screw microscope into the mouse’s baseplate until it is secured.
4. Place the mouse into a clean mouse cage and allow it to explore freely.

### Assess the field of view for cellular activity

5. Turn on the computer with the acquisition software and the DC power source which will serve as the power source for the microscope’s LED.
6. Plug the camera’s USB into the computer, and connect the microscope’s female BNC connector to the male BNC on the power source.
7. Start the acquisition software by entering in the Command Prompt: `> python “path/to/file/vid.py”` A new window displaying the camera’s live feed and default recording settings (20 fps, 15 gain) will launch.
8. Power the LED by increasing the voltage on the DC power source. Typical voltage settings range from 1.5-3.0 V.
9. If GCaMP-expressing neurons are visible within the field of view, adjust the voltage on the power source and/or the gain from the computer (by pressing “[“ or “]” for −1 and +1 gain, respectively) to determine the ideal illumination settings to use during the imaging experiment. *Apply the lowest possible voltage to the LED to bring GCaMP-expressing cells into view. Intense illumination from the LED may generate heat and risk bleaching GCaMP fluorescence. Ideal settings will result in images with low intensity background signals, high signal-to-noise in areas containing GCaMP-expressing neurons during cellular activity, and no over-exposed areas within the field of view. See Table 1 for potential problems that may prevent visualizing neurons.*
10. Record the voltage from the DC power source, camera gain, and microscope LED position (angle between mouse’s nose and LED) for later use throughout the imaging experiment.

### Habituate mice to microscope procedures

11. Remove one mouse from the home cage, attach the microscope to the baseplate, and place the mouse in a new cage.
12. Tether the microscope wires to a surface above the cage (e.g., shelf above the mouse’s cage), and allow mouse to explore the cage for 10-15 min. *Ensure that there is enough slack in the cabling for the mouse to freely explore the entire area of the cage.*
13. Detach microscope and return the mouse to the home cage.
14. Repeat habituation procedure for 3-5 days before the start of the imaging experiment to ensure that all mice are comfortable with attachment/detachment procedures and can carry the microscope for timescales similar to the upcoming imaging sessions without experiencing fatigue. *Mice with low body weights (less than 20 g) may require additional habituation sessions to comfortably carry the microscope.*

## Basic Protocol 5: *IN VIVO* imaging during fear behaviour

The following protocol describes use of the CHEndoscope system to acquire imaging data from an awake, behaving mouse during a Pavlovian fear conditioning experiment. The main feature of this protocol involves synchronization of the CHEndoscope acquisition software with the behavioural software controlling trial duration and timing of stimulus presentation. Here we detail use of FreezeFrame software from Coulbourn Instruments, but in principle, any software capable of sending commands through UDP can be used to synchronize signals between the behavioural apparatus and the CHEndoscope.

### Materials

- Adjustable DC power source
- FreezeFrame software (Colbourn Instruments, Allentown, PA, USA)
- Computer running FreezeFrame
- Computer running CHEndoscope acquisition software
- Fear chamber equipped with camera and stimulus (shock) generator
- 70% EtOH
- Clean mouse cage
- CHEndoscope

### Set up behavioural and recording apparatuses

1. Start FreezeFrame software and open a Command Prompt window for running the CHEndoscope acquisition software. *Important: If both pieces of software are being run on the same computer, ensure the computer is powerful enough to encode the necessary videos without dropping frames. Alternatively, run FreezeFrame and the CHEndoscope acquisition software on separate computers and connect them to a wired ethernet network*.
2. In the recording computer’s Command Prompt window, set the working directory to the location where recording files (.mkv video and .pkl file) will be saved: `> cd “Path/to/folder”`
3. On the behaviour computer, open FreezeFrame and set the mode to “Remote”, allowing communication between the FreezeFrame program and the acquisition software.
4. Under Settings>Remotes, turn on Remote 1, and in the Communication window, enter the IPv4 address from the recording computer in the box adjacent to Remote 1. *On Windows systems, the IPv4 address can be found under the “Ethernet Adapter” heading after running `>ipconfig` in the Command Prompt*.
5. Under Settings>Cameras, set the frame rate of the behavioural recording to 15 fps and restart the program to refresh the camera settings.
6. Set the working directory where the behavioural recordings (.ffii files) will be saved, and select or create the desired behavioural protocol including shock US stimuli for conditioning from the dropdown in FreezeFrame.
7. Set the shock intensity for the US on the stimulus generator (typically, between 0.4-0.7 mA).
8. Clean the inside of the fear chamber with 70% EtOH and dry completely with a paper towel.

### Record calcium activity with microendoscope during fear conditioning

9. Transfer mice to holding room approximately 1 h before first recording session.
10. Using a clean mouse cage, transfer one mouse to the recording room.
11. Attach CHEndoscope to the mouse’s baseplate, connect the CHEndoscope’s USB connector to the recording laptop and the BNC connector to the power source, and return the mouse to the cage.
12. Run recording software in vid.py by entering in Command Prompt: `> python “Path/to/file/vid.py”`
13. Ensure that the two programs are communicating by acquiring the “Reference” image in FreezeFrame. A line reading “[refc]” should appear immediately in the Command Prompt window if the programs are properly synchronized.
14. Check the field of view by powering the LED using the DC power source and setting the voltage on the power source and gain on the computer to the parameters determined previously (in Basic Protocol 4). Adjust gain (using “[“ and “]” keys to decrease and increase gain, respectively) and/or LED voltage if necessary and record settings used during the imaging session.
15. Place the mouse in the fear chamber, close the door, and quickly tether the microscope’s wires above the fear chamber so that the mouse is free to explore the entire area of the chamber.
16. In FreezeFrame, start the training protocol by pressing the Start button, which will also initiate the calcium imaging recording.
17. Wait until the end of the protocol, then remove the mouse from the fear chamber, detach the microscope, and return the mouse to the home cage.
18. Clean the chamber with 70% EtOH, and repeat steps 12-19 for all mice to acquire calcium imaging recordings during fear conditioning.

### Record calcium activity with CHEndoscope during fear recall

19. Repeat steps 1-18 on a subsequent day, with the exception of omitting shocks (i.e., context test with no additional US presentations) by selecting a different behavioural protocol in FreezeFrame.

## Alternate Protocol: *In vivo* imaging during open field exploration

In this alternate protocol, we describe use of the CHEndoscope to record neuronal activity during unrestrained spatial exploration in an open field environment. We include instructions for food restricting mice prior to the start of the experiment, which increases motivation to explore during exposure to the open field. Next, we describe how to record calcium activity simultaneously with spatial exploration behaviour using behavioural recording software. Notably, this protocol can be easily applied to behavioural experiments involving spatial exploration of different environments (e.g., conditioned place preference, linear track, T- or Y-maze tasks, etc.).

### Materials

- Scale (for weighing mice and food)
- Food rewards (sprinkles, sucrose pellets, or other)
- Petri dishes
- Adjustable DC power source
- Computer running CHEndoscope acquisition software
- Computer running behavioural recording software
- Camera for recording mouse behaviour
- Open field apparatus
- 70% ethanol
- Clean mouse cage
- CHEndoscope

### Food restrict mice

1. Weigh mice with *ad libitum* food access daily for 5 days to determine free-feeding weight (average weight across days).
2. At least one week prior to the start of the experiment, restrict food intake of mice by feeding small amounts of food daily to reduce body weight to 90% of their free-feeding weight. *Maintain mice at 90% body weight throughout the duration of the experiment by weighing and feeding mice daily.*
3. Expose mice to food rewards (e.g., sprinkles, sucrose pellets, etc.) daily for 3-5 days before the experiment by placing a petri dish containing 5-10 rewards per mouse in the home cage at feeding times. Monitor consumption to ensure that all mice eat the rewards. *Reward exposure can be performed on the same days as microscope habituation procedures (see Basic Protocol 4).*

### Set up behavioural and recording apparatuses

4. On the recording computer, open Command Prompt and set the working directory where the recording files (.mkv video and .pkl file) will be saved: `> cd “Path/to/folder”`
5. Mount camera for recording behaviour above the open field apparatus and start computer with the behavioural recording software (e.g., ANY-maze, Ethovision).
6. Set recording duration (typically, 10 min) by designing a protocol within the behavioural software.
7. Clean the bottom of the open field with 70% EtOH and dry with a paper towel.
8. Scatter food rewards (15-20 total, same as those used in step 4) over the entire area of the open field to increase motivation of mice to explore the entire field.

### Record calcium activity with microendoscope during exploration behaviour

9. Transfer mice to holding room approximately 1 h before the first recording session.
10. Using a clean mouse cage, remove one mouse and transfer to recording room.
11. Attach CHEndoscope to the mouse’s baseplate, connect the CHEndoscope’s USB to the recording laptop and the BNC connector to the power source, and return the mouse to the cage.
12. Run recording software by entering in Command Prompt: `> python “Path/to/file/vid.py”`
13. Check the field of view by powering the LED using the DC power source and setting the voltage on the power source and gain on the computer to the parameters determined previously (in Basic Protocol 4). Adjust gain (using “[“ and “]” keys to decrease and increase gain, respectively) and/or LED voltage if necessary and record settings used during the imaging session.
14. Place the mouse in the open field and start the behavioural recording and calcium imaging recording.
15. Allow the mouse to freely explore the open field for 10 min.
16. At the end of the session, remove the mouse from the open field, power off the LED, and remove the microscope before returning the mouse to its home cage.
17. Discard any food rewards and feces left by the mouse in the open field, clean the bottom of the chamber with 70% EtOH, and scatter new food rewards around the open field for the next mouse.
18. Repeat recording procedure (steps 12-19) for all mice, and on multiple days as required by the experimental design.

## Basic Protocol 6: Histology

Following every experiment, processing of brain tissue is necessary to verify the placement of the GRIN lens and if applicable, assess the AAV infection in the target tissue. With a few exceptions, this protocol follows typical methods for preparing brain tissue for immunohistochemistry assays. If desired by the experimenter, immunohistochemistry can be performed on the same brain tissue to identify cells expressing specific markers. Representative brain sections showing GCaMP expression and GRIN lens placements are shown in Figure 4.

**Figure 4.**
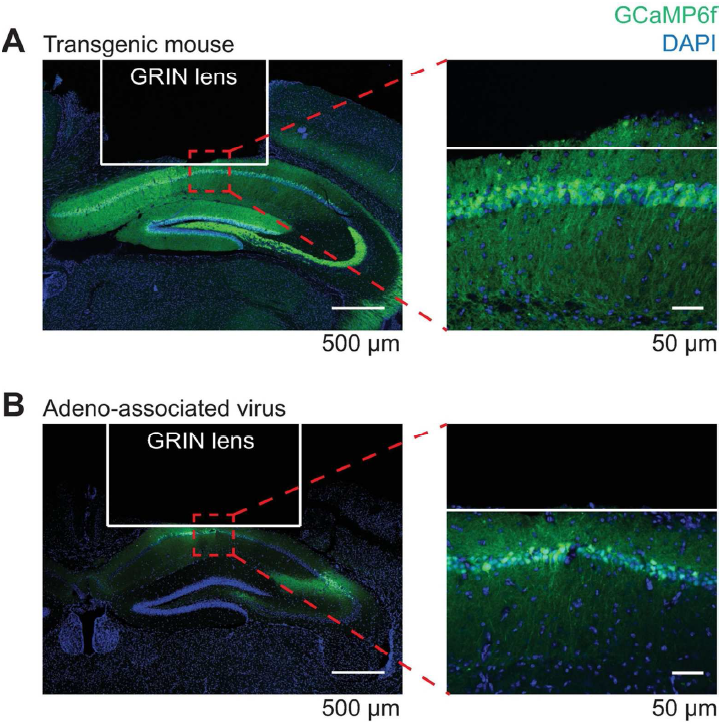
Histology. (A) Representative confocal image showing GRIN lens placements above CA1 from a transgenic mouse expressing GCaMP6f under the Thy-1 promoter. (B) GRIN lens placement from a wild-type mouse infused with an AAV carrying the GCaMP6f transgene under the CaMK2α promoter. 108x108mm (300 × 300 DPI)

### Materials

- Appropriate anesthetic
- Perfusion equipment
- 1x phosphate-buffered saline (1xPBS), chilled at 4 °C (see Reagents)
- 4% (w/v) paraformaldehyde (PFA), chilled at 4 °C (see Reagents)
- 15- and 50-ml Falcon tubes
- 30% (w/v) sucrose (see Reagents)
- 12- or 24-well plate
- Cryostat or vibratome
- Paintbrushes for sectioning and mounting tissue
- DAPI stock solution
- Gel-coated microscope slides
- Glass coverslips
- Anti-fading mounting medium
- Confocal microscope (for imaging slides)

### Perfuse mice

1. Deeply anesthetize mouse with appropriate anesthetic.
2. Expose the thoracic cavity by cutting through the diaphragm and extending the incision up both sides of the mouse along the rib cage.
3. Perfuse the mouse with 40 ml 1xPBS followed by 40 ml PFA.
4. Decapitate the mouse and submerge the head in 50-ml Falcon tube containing approximately 20 ml PFA.
5. Store the mouse heads for 2-5 overnights (maximum 2 overnights if performing immunohistochemistry) at 4 °C to allow the brains to post-fix around the GRIN lens. *Longer post-fixation periods result in cleaner lens tract in the tissue.*

### Remove and section brains

6. After post-fixation, remove the headcap from the skull by loosening the screws with fine scissors then pulling the headcap straight up with large forceps or hemostats. *Minimize wiggling the headcap during removal to avoid possible damage to the imaging surface or widening the lens tract.*
7. Carefully remove the brain from the skull and place in a 15-ml Falcon tube filled with 30% sucrose. Protect from light and store at 4 °C for 2-3 days until ready to section.
8. Section the brain into 50 um slices using a cryostat or vibratome, collecting the entire anterior-posterior extent of the GRIN lens tract.
9. Protect brain sections from light before staining.

### Stain and mount tissue

10. Perform immunohistochemistry if necessary, avoiding conjugates whose spectra overlap with GCaMP6f. *The GCaMP6f signal does not typically require amplification with antibodies.*
11. Wash sections for 3x 10 min in 1xPBS at room temperature (RT).
12. Counterstain tissue with DAPI (1:10,000 DAPI stock in 1xPBS; DAPI stock, 5 mg/ml in dH_2_O) for 10 min at RT.
13. Wash sections for 3x 10 min in 1xPBS at RT.
14. Mount sections on gel-coated slides and coverslip using an anti-fading medium (e.g., PermaFluor).

### Verify lens placements with microscopy

15. Image slides using a confocal microscope to verify that GRIN lens placements are above the target region, and that cells below the GRIN lens look healthy. *Imaging should be performed on sections where the lens tract is widest, indicating the center of the lens which has the greatest probability of containing the imaged cell population.*
16. If necessary, image additional sections to observe the extent of the viral infection.

## Basic Protocol 7: Analysis of Imaging Data

Videos obtained from the acquisition phase of the experiment must be processed in order to retrieve individual cell calcium transients (cell activity data). The CHEndoscope analysis workflow (Figure 5) involves three main steps. First, the raw videos are downsampled and motion-corrected. Next, using Constrained Nonnegative Matrix Factorization for microendoscopic data (CNMF-E; Zhou et al., 2017), traces and spatial footprints of putative neurons are extracted. These are manually inspected using custom code to remove traces that do not meet certain criteria. Lastly, neurons are matched between sessions using CellReg (Sheintuch et al., 2017). Data retrieved at the end of the analysis workflow can be used for experiment-specific analyses.

**Figure 5.**
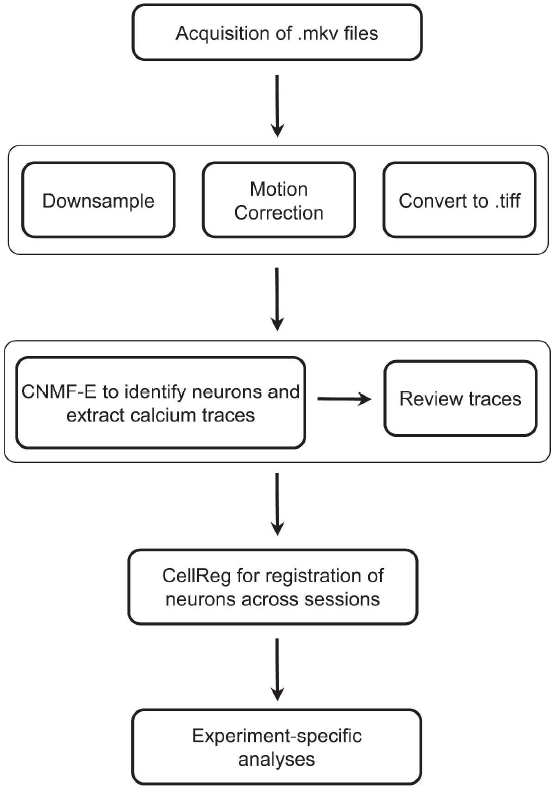
A schematic of the processing and analysis pipeline beginning after acquisition of imaging videos in preparation for experiment-specific analyses. (CNMF-E) Constrained Nonnegative Matrix Factorization for microendoscopic data. 559×783mm (300 × 300 DPI)

**Figure 6.**
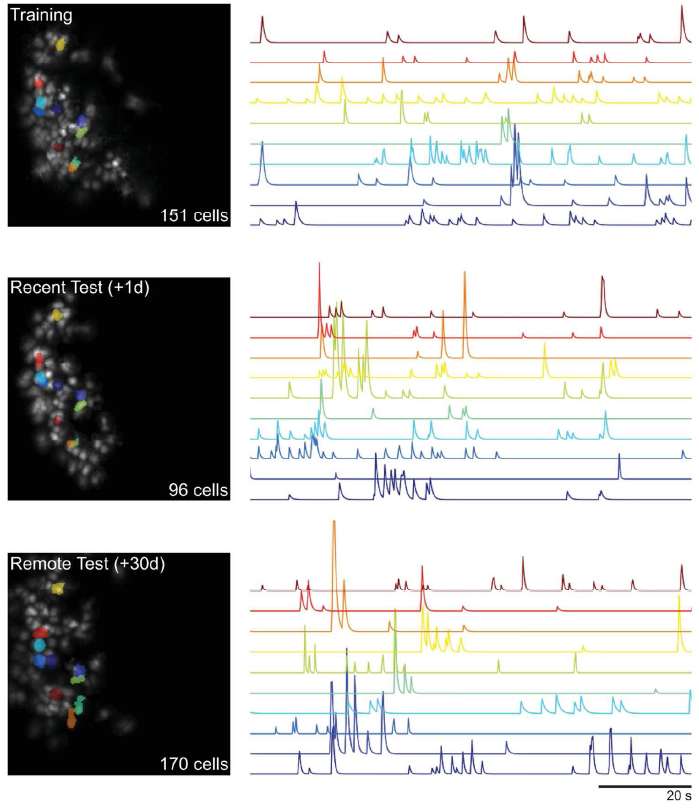
Examples of CA1 neurons from a month-long fear conditioning experiment. Spatial footprints of cells extracted from each session using CNMF-E are shown (left) with a segment of the denoised calcium trace for ten cells (right). Neurons that did not meet review criteria (see Basic Protocol 7) are not shown. Neurons were tracked across sessions using CellReg (spatial footprint and trace identity are indicated by colour). 132x152mm (300 × 300 DPI)

### Materials

- <u>Python 3.5+</u> including the following packages and versions: numpy ver. 1.12.0 scikit_image ver. 0.13.dev0 matplotlib ver. 1.5.3 scipy ver. 0.18.1 joblib ver. 0.11 pims ver. 0.4 skimage ver. 0.0 scikit_learn ver. 0.19b2 tqdm ver. 4.14.0
- <u>Mkvmerge:</u> merge mkv files
- <u>Tiffcp:</u> convert and merge .tiff files

### Preprocess videos

1. Change the directory to the location of the minipipe folder through the command line: `> cd “Path/to/jflab-minipipe/”`
2. Run minipipe.py with python on the videos to be downsampled and/or motion corrected. `> python minipipe.py /path/to/file.mkv` *If running into memory errors at this step, change chunk size to a smaller value, i.e., add the* command `flag -c size` *where size is your new chunk size (default chunk size is 2000 frames).*
3. Open the output .tiff video in ImageJ or FIJI to manually inspect the applied transformations from the motion correction.

### Identify Neurons and Extract Calcium Traces

4. Run CNMF-E as described at https://github.com/zhoupc/CNMF_E. *CHEndoscope-optimized parameters for CA1 imaging are listed in Table 2*.
5. Run the review_traces.py script on the motion corrected video. `> python review_traces.py /path/to/file.mat`
6. In the graphical user interface (GUI), choose to keep or exclude each identified neuron using the ‘keep’ and ‘exclude’ buttons, or the ‘j’ and ‘k’ keys for exclude and keep, respectively. *Inspect the extracted calcium transients and spatial footprints based on the previously reported criteria*:

**Table 2.**
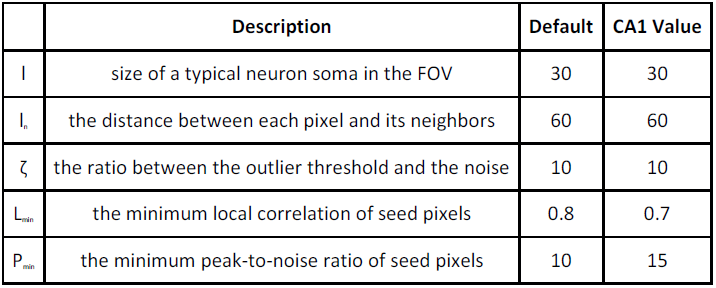
User-selected parameters for the CNMF-E algorithm and optimized values used for processing hippocampal CA1 videos from the CHEndoscope.

|  | Description | Default | CA1 Value |
| --- | --- | --- | --- |
| $l$ | size of a typical neuron soma in the FOV | 30 | 30 |
| $l_n$ | the distance between each pixel and its neighbors | 60 | 60 |
| $\zeta$ | the ratio between the outlier threshold and the noise | 10 | 10 |
| $L_{\min}$ | the minimum local correlation of seed pixels | 0.8 | 0.7 |
| $P_{\min}$ | the minimum peak-to-noise ratio of seed pixels | 10 | 15 |

a. *fast rise and slow decay of calcium transients with stable baseline fluorescence (Resendez et al., 2016)*.
b. *calcium transient durations consistent with GCaMP6f (or appropriate GCaMP variant) (Badura, Sun, Giovannucci, Lynch, & Wang, 2014)*.
c. *spatial footprint is consistent with shape and size of neuron (Resendez et al., 2016)*.

### Register Neurons Between Sessions

7. Run the reshape_spatial_footprints.py script on the output file from step 6 to format the file for use in CellReg. `> python reshape_spatial_footprints.py “/path/to/file.mat”`
8. Run CellReg as described at https://github.com/zivlab/CellReg.

## Reagents and Solutions

### Artificial cerebrospinal fluid (aCSF, 500 mL)

400 mL dH2O

3.65 g sodium chloride (0.365% w/v)

0.186 g potassium chloride (0.0186% w/v)

0.901 g D-(+)-Glucose (0.0901% w/v) 1.192 g HEPES (0.119% w/v)

0.246 g magnesium sulfate heptahydrate (0.0246% w/v)

0.147 g calcium chloride dihydrate (0.0147% w/v)

Mix well and adjust to pH of 7.3

Add dH2O up to 500 mL

Sterilize by passing through a 0.2 um-pore filter

Store up to 1 month at 4ºC

### Phosphate-buffered saline (1x PBS, 1 L)

750 mL dH2O

8.52 g sodium phosphate dibasic anhydrous (0.852% w/v)

5.52 g sodium phosphate monobasic monohydrate (0.552% w/v)

9.0 g sodium chloride (0.9%)

Mix well and adjust to pH of 7.3

Add dH2O up to 1 L

Store up to 1 year at room temperature

### 16% paraformaldehyde (250 mL)

Bring 250 mL of dH_2_O to 50ºC

40 g paraformaldehyde (16% w/v)

Bring temperature to 60ºC

Add 0.5 mL 10.0 N NaOH (0.2% v/v)

Once PFA is dissolved, cool and filter

Store up to 2 weeks at 4ºC

*Do not exceed 65ºC to avoid degrading formaldehyde*

### 4% paraformaldehyde (400 mL)

100 mL of 16% paraformaldehyde (25% v/v)

300 mL of 1x PBS (75% v/v)

Store up to 2 weeks at 4ºC

## Commentary

### Background information

The central advantages of the CHEndoscope system are its simple construction, low cost, and customizable design. A complete CHEndoscope, including filter box, camera body and baseplate can be assembled in less than 6 hours without the use of specialized tools. Constructing a CHEndoscope does not require a large initial investment, as most of the necessary components are available at low cost from consumer electronics vendors. The surgical and behavioural equipment needed for the CHEndoscope system are similarly minimal; most laboratories outfitted for behavioural neuroscience research already own the specialized equipment necessary to perform the surgeries and behavioural assays outlined in this Unit. Furthermore, analysis of calcium imaging data described in Basic Protocol 7 can be performed using open-source code provided by our group and others (i.e., CNMF-E and CellReg).

In comparison to commercially available systems, the CHEndoscope offers the additional advantage of being customizable for specific applications. It is possible to modify the custom-printed housing of the CHEndoscope, allowing it to be adapted for use with electrophysiological stimulation and recording rigs, optogenetics, or fibre photometry. This modification process can be done using a variety of computer-aided design (CAD) software packages to modify the design files provided in our online repository. Once new designs are created, custom housings can be manufactured by a 3D printer within hours. Given the flexibility of the CHEndoscope’s design and its low cost, it is possible and practical to tailor the system to the particular needs of a research question.

### Critical parameters

While planning the experiment, careful consideration should be given to whether GCaMP expression in the target brain region will be achieved using local AAV infection or transgenic mice. The choice of using a viral or transgenic approach to express GCaMP will depend on the aim and design of the specific imaging experiment. Our laboratory has successfully recorded calcium activity from CA1 pyramidal neurons in both WT mice infused with AAVs encoding GCaMP6f (DJ-serotype, hsyn or CaMK2α promoters) and transgenic GP5.17 mice (Jackson Laboratories, Stock No. 025393, RRID: IMSR_JAX:025393) which show robust expression of GCaMP6f under the Thy-1 promoter in the hippocampus and cortex (Dana et al., 2014) (see Figure 4). In our hands, these methods have produced similar results in fear conditioning experiments. However, caveats exist for both viral and transgenic approaches, and the limitations of each method may be unmasked by different experimental designs. For example, we generally recommend using transgenic mice for calcium imaging experiments involving repeated, chronic recordings (i.e., recordings over multiple weeks to months), since GCaMP expression in transgenic lines is stable through a majority of the animals’ lifespan (up to 11 months of age, and possibly longer), reducing the risk of overexpression, cytotoxicity, and cell death over time (Dana et al., 2014; Madisen et al., 2015). Importantly, while transgenic GCaMP mice may be more suitable for longterm imaging, researchers should be aware that multiple GCaMP6 mouse lines have been shown to exhibit aberrant, epileptic-like cellular activity - possibly caused by early developmental expression of GCaMP6 and/or calcium buffering in specific cell types (Steinmetz et al., 2017) - and thus the choice of transgenic mouse line should be made carefully.

In other situations it may be necessary to express GCaMP using viral methods, for example if transgenic mice show poor expression in the target brain region. Further, experiments that seek to image activity from subpopulations of neurons that are genetically-defined (Kamigaki & Dan, 2017) or defined by projection pattern (Murugan et al., 2017; Zhou et al., 2017) may require the use of Cre-dependent AAVs in combination with the appropriate Cre-expressing mouse lines or viruses. In any experiment relying on AAV-mediated GCaMP expression, experimenters should give adequate time between AAV infusion and imaging to allow expression of the calcium indicator to peak (typically, at least 1 month) and as mentioned above, ensure that the specific methods being used results in stable protein expression over the full duration of the imaging experiment (Resendez et al., 2016).

### Troubleshooting

**Table 1:** Troubleshooting guide for assessing the field of view (FOV) of cellular activity after recovery from lens implantation. Problems for which the mouse is no longer usable are noted as having no solution (N.S.).

| Problem | Possible Cause | Solution |
| --- | --- | --- |
| Inconsistent illumination of cells caused by flickering LED | Loose connection between filter box and LED power cable | Determine whether the connection between the LED power cable and two-pin female receptacle or soldered connection between two-pin female receptacle and LED is loose, and replace appropriate parts |
| Whole FOV shaking or moving | Loose connection between filter box and baseplate | Check baseplate ring for debris, clean out using size 5 forceps (ensure connection between baseplate and filter box is stable after reattaching the CHEndoscope) |
| Brightest spot in FOV shaking or moving | Dichroic mirror moving inside filter box | Open the filter box by removing the plastic cover, re-glue the dichroic mirror to the filter box, and re-seal the filter box with a new plastic cover |
| No cells visible with indistinct landmarks (e.g., blood vessels) | Insufficient recovery time | Allow mice to recover for an additional 1-3 weeks and re-check with the CHEndoscope |
| No cells visible with distinct landmarks (e.g., blood vessels) | Off-target virus injection and/or lens placement | N.S. |
| Few or blurry cells | Insufficient recovery time OR out-of-focus image | Allow mice to recover for an additional 1-3 weeks and re-check with the CHEndoscope OR change focal plane by loosening the camera body's nylon set screw and moving the camera body up or down to bring cells into focus |
| Extremely bright | Cell death due to | N.S. |
| cells with static signal | GCaMP overexpression or tissue damage |  |
| Dark patches or puncta (static or flowing) covering cells | Occlusion of tissue by blood clots or active bleeding | Allow 1-3 weeks for blood to clear and re-check with the CHEndoscope |
| Air bubbles between lens and cells | Bacterial infection caused by non-sterile surgical techniques | N.S. |

### Statistical analyses

Videos acquired during imaging sessions must go through a number of processing steps before analysis can begin. Raw videos are first temporally downsampled using the mean pixel values, reducing the file size in order to speed up the following processing steps. Motion artifacts within each video are corrected by aligning each video frame to a template frame (typically, the first frame). To do this, a low-pass spatial filter is applied to remove high frequency neuronal signals from each frame and enhance lower frequency structural features. These features are used to determine the values needed to align each video frame to the template frame.

After downsampling and motion correction, the output video file is run through CNMF-E, which outperforms previous algorithms such as PCA/ICA (Mukamel, Nimmerjahn, & Schnitzer, 2009). Following CNMF-E, the relevant data are saved as a .mat file and traces are manually reviewed using custom code written in Python. Manual inspection of neurons detected by CNMF-E is required to remove false positives. The review traces program displays the raw and denoised traces, along with the spatial footprints of putative neurons, and gives users the ability to choose which neurons to exclude from further analyses. Finally, for experiments in which there are multiple recording sessions either within or across days, CellReg (Sheintuch et al., 2017) is used to identify the same cells across sessions. CellReg uses a probabilistic model to determine whether spatial footprints from two sessions align with each other.

Particular statistical analyses performed on cell activity data will depend on experiment-specific hypotheses and questions. The statistical analyses used for neuron/trace detection in CNMF-E have been previously described (Pnevmatikakis et al., 2016; Zhou et al., 2017)). Tracking of cells between sessions in CellReg relies on a probabilistic statistical model of spatial footprint matching (Sheintuch et al., 2017).

### Understanding results

The protocols in this unit describe the collection and extraction of calcium activity traces from a population of neurons during behaviour. This constitutes a pre-processing stage, and further analyses are required to examine the relationship between neuronal activity and behaviour. For example, properties of individual neurons can be identified using calcium transient data. Previous studies have identified position-encoding neurons in the hippocampus and cortex (Murugan et al., 2017; Wagatsuma et al., 2018) and stimulus or expectation-encoding neurons in the amygdala (Yu et al., 2017). Analysis at the population level has yielded insights into the representations of social stimuli (Li et al., 2017; Yu et al., 2017) and fearful memories in the amygdala (Grewe et al., 2017).

### Time considerations

Assembly of the CHEndoscope filter box and camera body can be completed in approximately 6 hours spread across 2 days. While filter boxes and camera bodies need to be constructed only infrequently, one baseplate must be assembled for each implant surgery. Baseplate construction can be completed in less than an hour, with an additional 1.5 days required for the two glue-drying steps. Large numbers of baseplates can be assembled in batches for increased efficiency.

As noted in Basic Protocol 4, recovery after implantation of the baseplate and GRIN lens typically requires 3-6 weeks before clear, GCaMP-expressing neurons can be visualized using the CHEndoscope. In addition, habituation procedures before the start of behaviour and imaging may require up to 1 week. Timelines for specific calcium imaging experiments will vary depending on the specifics of the experimenter’s behavioural protocol.

## Acknowledgements

We would like to thank Valentina Mercaldo, Ying Meng, Tiange Li and Yasaman Soudagar for early design input; Antonietta De Cristofaro, Daisy Lin, and Mika Yamamoto for excellent technical assistance; and Mazen Kheirbek (UC San Francisco) and Jessica Jimémez (Columbia University), and Hendrik Steenland (NeuroTek Innovative Technology Inc.) for collaboration. In addition, we would like to acknowledge training from Canadian Neurophotonics Platform. This work was supported by grants from the Canadian Institutes of Health Research (S.A.J, P.W.F, C.Y), Natural Sciences and Engineering Research Council of Canada (S.A.J., P.W.F., A.D.J., A.I.R., A.J.M., L.M.T., C.Y), Canadian Institute For Advanced Research (S.A.J., P.W.F.). In addition, A.J.M., L.M.T. and C.Y. received support from Restracomp Fellowships (Hospital for Sick Children).

